# Abrupt Pericyte Loss Precedes Endothelial Activation in Cerebral Small Vessel Disease

**DOI:** 10.64898/2026.03.19.713028

**Authors:** A. Chagnot, D. Jaime Garcia, C. McQuaid, J. Cholewa-Waclaw, K. McDade, O. Dando, J.M. Wardlaw, C. Smith, A. Montagne

**Affiliations:** Institute for Neuroscience and Cardiovascular Research (INCR), Chancellor’s Building, The University of Edinburgh, 49 Little France Crescent, Edinburgh EH16 4SB, UK; Centre for Clinical Brain Sciences (CCBS), School of Clinical Sciences, College of Medicine and Veterinary Medicine, The University of Edinburgh, Edinburgh EH16 4SB, UK; UK Dementia Research Institute (UK DRI) at the University of Edinburgh, Edinburgh EH16 4SB, UK; British Heart Foundation - UK Dementia Research Institute (BHF-UK DRI) Centre for Vascular Dementia Research (CVDR) at the University of Edinburgh, Edinburgh EH16 4SB, UK; Centre for Regenerative Medicine (CRM), Institute for Regeneration and Repair (IRR), The University of Edinburgh, Edinburgh EH16 4UU, UK; Simons Initiative for the Developing Brain (SIDB), The University of Edinburgh, Edinburgh EH8 9XF, UK

**Author notes:** Contributed equally (A. Chagnot, D. Jaime Garcia, C. McQuaid).

**Keywords:** Cerebral small vessel disease, pericytes, capillaries, ageing, endothelial activation

## Abstract

Cerebral small vessel disease (cSVD) is a leading cause of stroke and vascular cognitive impairment, yet the cellular mechanisms underlying microvascular dysfunction in human disease remain incompletely understood. In particular, the relationship between pericyte alterations and endothelial activation, two key features of blood-brain barrier dysfunction, remains unresolved. Here, we performed a quantitative single-vessel analysis of the human cortical microvasculature across increasing cSVD severity and ageing. Using multiplex immunohistochemistry combined with spectral unmixing and automated image analysis, we analysed 11,409 microvascular fragments from post-mortem brain tissue derived from 20 cases. Endothelial cells, pericytes, and endothelial activation were assessed using PECAM-1, PDGFRβ, and VCAM-1, respectively. Microvascular density and diameter differed between cortical grey matter and the underlying white matter, with white matter vessels being less dense and wider in controls. While vessel diameter remained stable across disease stages, microvascular density increased with cSVD severity and age in the white matter. At the molecular level, PDGFRβ signal decreased markedly with increasing cSVD severity, consistent with progressive pericyte loss. This reduction was observed in both grey and white matter and correlated with disease severity and age. Notably, intermediate disease groups displayed marked heterogeneity, with vessels exhibiting either preserved or near-complete pericyte coverage, suggesting a potentially bimodal transition. In parallel, endothelial markers PECAM-1 and VCAM-1 increased significantly with disease severity, reflecting endothelial activation. Unsupervised Gaussian mixture clustering of marker expression identified three vascular states characterised by (i) preserved pericytes with low endothelial activation, (ii) marked pericyte loss without endothelial activation, and (iii) combined pericyte depletion and endothelial activation. These clusters broadly aligned with clinical severity but revealed intermediate states not captured by post-mortem diagnosis alone. Together, these findings suggest that pericyte loss and endothelial activation are partially dissociated processes that occur in a sequential progression in human cSVD, supporting pericyte dysfunction as an early event and highlighting it as a potential therapeutic target in microvascular disease.

## Introduction

Cerebral small vessel disease (cSVD) is a leading cause of stroke and the principal vascular contributor to cognitive impairment and dementia. It affects the brain’s perforating arterioles, capillaries, and venules, and is characterised by a spectrum of radiological and pathological features including white matter hyperintensities (WMH), lacunes, microbleeds, and blood-brain barrier (BBB) dysfunction[52]. Despite its high prevalence and clinical impact, there are currently no disease-modifying therapies, although recent clinical trials targeting cSVD mechanisms have shown promise, highlighting an incomplete understanding of the cellular mechanisms that initiate and propagate microvascular injury[34, 53].

At the capillary level, the cerebral microvasculature forms a key part of the neurogliovascular unit (NGVU), a multicellular system in which endothelial cells and pericytes maintain BBB integrity and regulate vascular function. Endothelial cells form the structural basis of the BBB and regulate molecular transport, immune surveillance, and vascular tone. Pericytes, embedded within the capillary basement membrane, provide structural support, regulate capillary diameter to control blood flow, and support endothelial stability through platelet-derived growth factor-B/platelet-derived growth factor receptor-β (PDGF-B/PDGFRβ) signalling[2, 12]. Disruption of endothelial-pericyte crosstalk is increasingly recognised as a central feature of microvascular dysfunction in ageing and neurodegenerative disease[1, 9, 42].

Endothelial activation is a consistent hallmark of cSVD and represents a shift towards a pro-inflammatory and pro-thrombotic phenotype. Upregulation of adhesion molecules such as vascular cell adhesion molecule-1 (VCAM-1), intercellular adhesion molecule-1 (ICAM-1), and selectins promotes leukocyte adhesion, vascular inflammation, and BBB disruption[23, 48, 52]. Neuroimaging studies further demonstrate associations between endothelial activation and impaired cerebrovascular reactivity, reduced cerebral blood flow (CBF), and increased BBB permeability[7, 25, 49]. Circulating biomarkers of endothelial activation have also been linked to cSVD burden and progression, although findings remain heterogeneous and mechanistic interpretation is limited by the lack of spatial resolution[24]. For example, soluble VCAM-1 has been identified as a circulating mediator of ageing-associated cognitive decline, linking vascular inflammation to brain dysfunction[54]. More recently, large-scale proteomic studies have revealed both shared and distinct molecular signatures across different radiological manifestations of cSVD, highlighting the biological heterogeneity of the disease and the involvement of multiple vascular wall compartments[20]. In parallel, biofluid studies provide further evidence of pericyte injury in ageing and Alzheimer’s disease (AD), with increased levels of soluble PDGFRβ in cerebrospinal fluid associated with BBB dysfunction, vascular damage, and cognitive decline[35, 37, 38].

Pericyte dysfunction has also emerged as a key contributor to microvascular pathology. Experimental models consistently demonstrate that pericyte loss leads to reduced CBF, endothelial destabilisation, and BBB breakdown[4, 17]. Mechanistically, disruption of pericyte-endothelial signalling induces a pro-inflammatory endothelial phenotype, including upregulation of VCAM-1 and ICAM-1, increased leukocyte adhesion, and altered vascular homeostasis[32, 39, 50]. These findings support a model in which pericytes actively restrain endothelial activation under physiological conditions.

However, whether pericyte loss is a consistent and defining feature of human ageing and cSVD remains unresolved. Several neuropathological studies report reduced pericyte coverage and microvascular alterations in AD and other dementias, particularly in white matter regions[14, 19, 44]. In contrast, other studies report preserved pericyte coverage, or even evidence of microvascular remodelling without overt pericyte loss in AD[16, 36]. Recent transcriptomic analyses further highlight marked heterogeneity in vascular cell states across sporadic and familial AD, suggesting that pericyte alterations are context-dependent and may vary across disease stage, vascular compartment, and pathological subtype rather than representing a uniform loss[45].

These discrepancies highlight a central unresolved question: whether pericyte alterations represent a primary driver of microvascular dysfunction, a secondary consequence of endothelial injury, or a dynamic process evolving across disease progression. Importantly, many existing studies rely on bulk or region-averaged analyses, which may obscure heterogeneity at the level of individual microvessels and contribute to conflicting findings.

Consistent with this, cSVD is increasingly recognised as a disorder of microvascular function and cellular composition rather than a disease primarily defined by remodelling of larger vascular branches such as arteries and veins. Structural features such as vessel density and diameter are often relatively preserved, particularly in early disease, suggesting that cellular and molecular alterations within intact vascular networks are key drivers of pathology[41, 42]. Recent advances in multiplex imaging and quantitative pathology now enable detailed interrogation of the cerebral microvasculature at single-vessel resolution. These approaches allow simultaneous assessment of endothelial and perivascular markers, providing an opportunity to resolve the spatial and temporal relationship between pericyte alterations and endothelial activation across disease stages.

In this study, we test the hypothesis that pericyte alterations and endothelial activation represent temporally and spatially distinct, yet mechanistically linked processes that together may explain key aspects of microvascular pathology in cSVD. By applying high-resolution quantitative analyses to human brain tissue spanning the spectrum of disease severity, we aim to define the extent and distribution of pericyte alterations, characterise endothelial activation, and determine how these processes relate within the brain microvasculature.

## Methods

### Human tissue acquisition, study cases, and clinical assessment

Brain sections in Brodmann area 4 including the premotor cortex and the underlying white matter were obtained from 20 cases. The tissue originated from three large Scottish cohorts: the Lothian Birth Cohort (LBC)[13], the Lothian study of Intracerebral Haemorrhage, Pathology, Imaging and Neurological outcome (LINCHPIN)[43], and the Edinburgh Sudden Death Brain Bank[47]. The Scotland A Research Ethics Committee approved this study (10/MRE00/23). The Edinburgh Sudden Death Brain Bank is funded by the UK Dementia Research Institute and was established in 2003 to increase access to younger control cases. All sudden deaths require a post-mortem examination by the Procurator Fiscal and Forensic Pathology Department who identify suitable cases of presumed healthy brains; most sudden death cases are sudden cardiac deaths. The Edinburgh Brain Bank functions within the Human Tissue (Scotland) Act of 2006 and has full ethical approval for the use of brain tissue in research (East of Scotland Research Ethics Service, ref 16/ES/0084). Informed consent was obtained from all participants in the LBC and LINCHPIN studies, and post-mortem authorisation was obtained from next of kin in all cases.

Relevant clinical information, demographics, and clinical history were collected by a trained research nurse. In sudden death cases this was extracted retrospectively from GP or hospital records. In LBC36 and LINCHPIN, the data were collected prospectively as part of the original LBC and LINCHPIN studies. Data included participant age at death, sex, prior diagnoses of hypertension, diabetes, hypercholesterolemia, heart disease, or atrial fibrillation, confirmed cause of death, post-mortem interval, and any relevant past medical history, including chronic disease, malignancy, stroke, inflammatory or rheumatologic disease, psychiatric illness, or cardiovascular disease (**Table 1**). Exclusion criteria included a post-mortem interval greater than five days and a known diagnosis of neurological or neurodegenerative disease other than cSVD.

**Table 1.** Sample characteristics. Brain tissue from 20 donors was included in this study. Age and post-mortem interval are reported as mean ± standard deviation (SD) [median]. Other values are reported as n (%). cSVD, cerebral small vessel disease. Group-wise significance was assessed using Kruskall-Wallis (a) or Fisher’s exact (b) tests. *, one missing value; **, three missing values.

|  | No cSVD Control | Mild cSVD | Moderate cSVD | Severe cSVD | Significance |
| --- | --- | --- | --- | --- | --- |
| N | 5 | 5 | 5 | 5 |  |
| Age at death (years) | 32.6 ± 8.1 [34] | 59.2 ± 18.4 [57] | 75.4 ± 7.7 [73] | 83.2 ± 8.8 [82] | **$p = 0.0043^a$** |
| Female sex | 2 (40.0) | 1 (20.0) | 1 (20.0) | 1 (20.0) | $p = 1^b$ |
| Post-mortem interval (h) | 89.0 ± 30.3 [99] | 90.2 ± 38.7 [103] | 59.6 ± 33.9 [74] | 61.2 ± 36.9 [49] | $p = 0.30^a$ |
| Hypertension | 0 (0.0) | 1 (20.0) | 3 (60.0) | 5 (100.0) | **$p = 0.0075^b$** |
| Hypercholesterolemia | 0 (0.0) | 1 (20.0) | 0 (0.0) | 1 (20.0) | $p = 1^b$ |
| Diabetes | 0 (0.0) | 1 (20.0) | 2 (40.0) | 0 (0.0) | $p = 0.56^b$ |
| Smoking | 2 (40.0) | 3 (75.0) * | 4 (80.0) | 0 (0.0) ** | $p = 0.32^b$ |
| Heart disease | 2 (40.0) | 2 (40.0) | 3 (60.0) | 1 (20.0) | $p = 0.92^b$ |

### Tissue preparation

Brain tissue processing is described in the respective LBC[13], LINCHPIN[43], and Edinburgh Sudden Death Brain Bank[47] protocols. Paraffin-embedded blocks of brain tissue were fixed using unbuffered 10% formalin for 48-72 hours prior to histological processing. Formalin-fixed, paraffin-embedded sections (4 μm thick, ∼2 cm^2^) in the primary motor cortex / precentral gyrus (Brodmann area 4) were selected to include both grey and white matter.

### cSVD assessment

There are no agreed, validated protocols for cSVD assessment. Neuropathological examination[21, 46] was carried out by an experienced neuropathologist (Prof. Colin Smith), blinded to clinical, radiographic, and genetic data. Cerebral small vessel disease (cSVD) burden, graded as no cSVD control (n=5), mild cSVD (n=5), moderate cSVD (n=5), or severe cSVD (n=5), was based on the presence and severity of vessel wall sclerosis (collagen deposition with loss of smooth muscle cells), increased vessel wall thickness, reduced lumen diameter, enlarged perivascular spaces, perivascular haemosiderin deposition, white matter rarefaction, lacunar infarcts, and microhaemorrhages[22, 46]. Two cases in the severe cSVD group also showed mild cerebral amyloid angiopathy on neuropathological examination.

### Immunohistochemistry

Experimenters were blinded to cSVD severity score, cohort of origin and all in vivo data. Immunohistochemistry was performed using the BOND RX Fully Automated Research Stainer (Leica Biosystems). Tissue sections mounted on slides were covered with BOND Universal Covertile (Leica Biosystems S21.4611) and processed using a Polymer Detection Kit (Leica Biosystems DS9800) and an HRP PLEX Kit (Leica Biosystems DS9914), which includes a base platform comprising peroxide block, post-primary reagent, polymer reagent, DAB chromogen, and haematoxylin counterstain, as well as the required chromogens green (Leica Biosystems DC9913) and red (Leica Biosystems DS9390).

Primary antibodies were applied following antigen retrieval using BOND Epitope Retrieval Solution 1 (Leica Biosystems AR9961). For endothelial cells, PECAM-1 (Abcam ab134168, 1:250 dilution; 20 min retrieval); for the pericytes, PDGFRβ (Abcam ab69506, 1:250 dilution; 20 min retrieval); and for endothelial activation, VCAM-1 (Abcam, ab134047, 1:500 dilution; 30 min retrieval) were used. A total of 150 μL of antibody solution in Leica antibody diluent (Leica Biosystems AR9352) was applied per slide, after which the instrument executed the sequential multiplex staining protocol.

### Multispectral imaging and microscopy

Stained tissue sections were imaged at 20× magnification using the PhenoImager HT Automated Quantitative Pathology Imaging System (Akoya Biosciences). For each individual case, eight to ten fields of view (929.74 × 697.31 μm area, 0.5 × 0.5 μm in-plane resolution), equally distributed between white and grey matter, were selected for multispectral imaging and spectral unmixing. Briefly, light from brightfield images was unmixed using a liquid-crystal tuneable band-pass filter (420-800 nm) and compared with the spectral signatures of the individual chromogens. An in-house spectral unmixing algorithm was then applied to isolate individual stain signals, yielding separate pseudo-fluorescent channels, analogous to fluorescence microscopy[31].

### Vascular segmentation

Microvascular features were extracted using a brightfield-based multiplex immunohistochemistry and quantitative image analysis pipeline (**Figure 1**). Grey and white matter regions were defined in the primary motor cortex (Brodmann area 4) (**Figure 1A**). Representative fields of view and individual vessel close-ups illustrate the distinct microvascular architecture in grey and white matter (**Figure 1B, C**). Chromogenic signals corresponding to PECAM-1 (DAB), PDGFRβ (VectorRed), and VCAM-1 (MethylGreen) were separated into distinct pseudo-fluorescent channels by spectral unmixing (**Figure 1D, E**). Blood vessels were segmented based on the PECAM-1 signal using Morph-O-Matic[10], a MATLAB-based (MATLAB R2021b, The MathWorks Inc.) tool (**Figure 1F, G**). The PECAM-1 signal was processed using the following steps: thresholding above the 9^th^ decile of image intensity, removal of particles smaller than 125 μm^2^, 1.5 μm dilation, 7.5 μm median smoothing, filling of holes smaller than 750 μm^2^, 1 μm erosion, and a final 2.5 μm median smoothing. Eight-connected logic was used to separate the resulting vascular objects, which were subsequently analysed independently. To reduce bias in signal quantification, PDGFRβ and VCAM-1 coverage, thresholded above their respective mean signal intensities, was calculated within PECAM-1-positive vascular objects. Metrics extracted from each vascular object included length, vessel-wide diameter, area, PECAM-1 signal intensity, PDGFRβ signal intensity, and VCAM-1 signal intensity.

**Figure 1.**
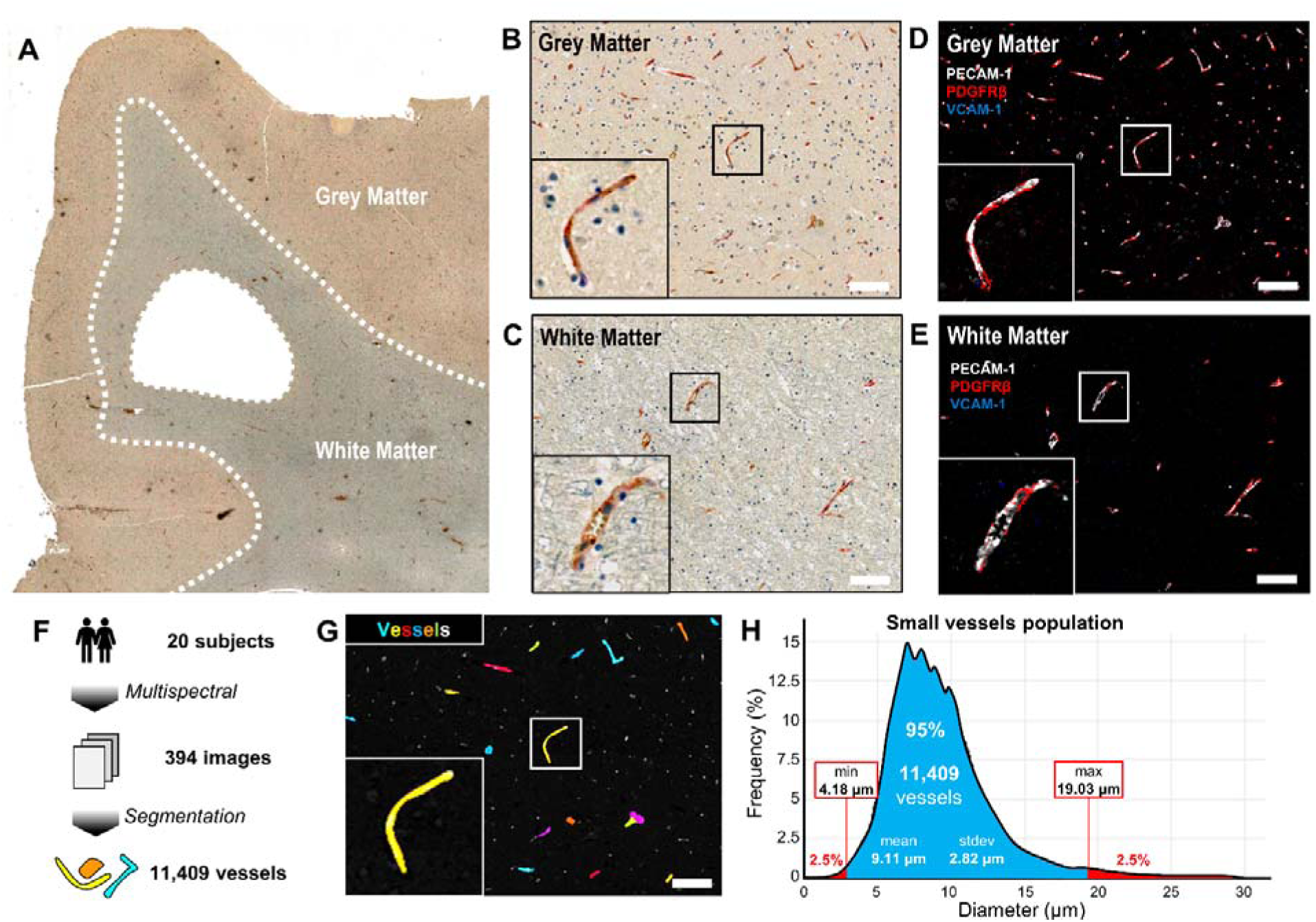
Extraction of microvascular features from multiplex brightfield imaging. (**A**) Representative brightfield image of a cortical brain section (Brodmann area 4, primary motor cortex) from a 34-year-old male no cSVD control, with annotated grey and white matter sampling regions. A minimum of eight fields of view (930 x 700 μm) were analysed per case. (**B**, **C**) Representative brightfield fields of view and corresponding vessel close-ups in grey (B) and white (C) matter. (**D**, **E**). Pseudo-fluorescent reconstructions of the images in B and C showing separated chromogenic signals for PECAM-1 (DAB), PDGFRβ (VectorRed), and VCAM-1 (MethylGreen). (**F**) Overview of the image processing and vascular segmentation workflow implemented in the MATLAB-based Morph-O-Matic[10] pipeline. (**G**) Segmentation of individual vascular fragments based on the PECAM-1 signal. (**H**) Diameter distribution of segmented vascular fragments; extreme values were excluded to remove segmentation artefacts. Scale bars: 200 μm. Abbreviations: DAB, 3,3’-diaminobenzidine; PECAM-1, platelet endothelial cell adhesion molecule-1; PDGFRβ, platelet-derived growth factor receptor-β; VCAM-1, vascular cell adhesion molecule-1.

The dataset was refined by excluding extreme vessel diameter values to remove segmentation artefacts smaller than the 2.5^th^ quantile (4.18 μm) and larger than the 97.5^th^ quantile (19.03 μm) leaving a total pool of vessels of 11,409 individual objects (mean diameter 9.11 μm, standard deviation 2.82 μm) from 20 cases (**Figure 1H**). For each of the 394 (8 to 10 per case and per matter, white or grey) fields of view, vascular density, averaged vascular diameters, average PECAM-1 signal intensity, average PDGFRβ signal intensity, and average VCAM-1 signal intensity were computed. All the averages were weighted by the length of individual vascular objects.

### Statistical analyses

Processing of the data was done using R version 4.2.0 (The R Core Team, 2022). Vessels with extreme diameters were excluded to minimise the inclusion of segmentation artefacts. Kruskall-Wallis or Fisher’s exact tests were computed on the epidemiological data presented in **Table 1** to investigate inter-group differences. Mean, standard deviation, and median of each metric were reported for each disease group and matter in **Table 2**. Log transform was applied to the vascular density and signal data to adjust to normal distribution. Twenty-seven outliers (of 394 images) were identified using Mahalanobis distance (p=0.05) across all five metrics and excluded.

**Table 2.**
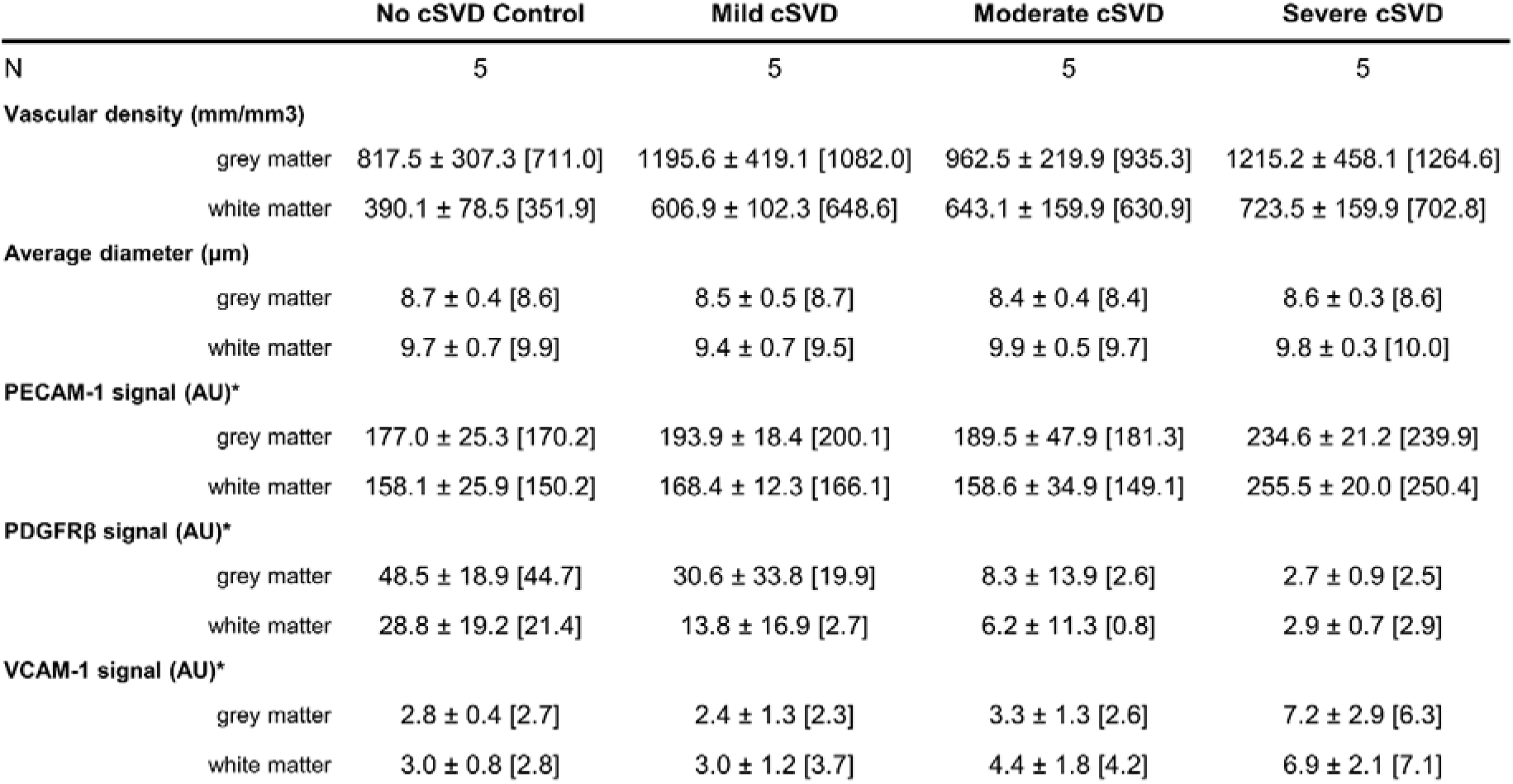
Quantitative vascular metrics by diagnostic group. All metrics are reported as mean ± standard deviation (SD) [median]. Abbreviations: AU, arbitrary units; cSVD, cerebral small vessel disease; PECAM-1, platelet endothelial cell adhesion molecule-1; PDGFRβ, platelet-derived growth factor receptor-β; VCAM-1, vascular cell adhesion molecule-1. *, values multiplied by 10^3^ for display.

|  | No cSVD Control | Mild cSVD | Moderate cSVD | Severe cSVD |
| --- | --- | --- | --- | --- |
| N | 5 | 5 | 5 | 5 |
| **Vascular density (mm/mm<sup>3</sup>)** |  |  |  |  |
| grey matter | 817.5 ± 307.3 [711.0] | 1195.6 ± 419.1 [1082.0] | 962.5 ± 219.9 [935.3] | 1215.2 ± 458.1 [1264.6] |
| white matter | 390.1 ± 78.5 [351.9] | 606.9 ± 102.3 [648.6] | 643.1 ± 159.9 [630.9] | 723.5 ± 159.9 [702.8] |
| **Average diameter (μm)** |  |  |  |  |
| grey matter | 8.7 ± 0.4 [8.6] | 8.5 ± 0.5 [8.7] | 8.4 ± 0.4 [8.4] | 8.6 ± 0.3 [8.6] |
| white matter | 9.7 ± 0.7 [9.9] | 9.4 ± 0.7 [9.5] | 9.9 ± 0.5 [9.7] | 9.8 ± 0.3 [10.0] |
| **PECAM-1 signal (AU)\*** |  |  |  |  |
| grey matter | 177.0 ± 25.3 [170.2] | 193.9 ± 18.4 [200.1] | 189.5 ± 47.9 [181.3] | 234.6 ± 21.2 [239.9] |
| white matter | 158.1 ± 25.9 [150.2] | 168.4 ± 12.3 [166.1] | 158.6 ± 34.9 [149.1] | 255.5 ± 20.0 [250.4] |
| **PDGFRβ signal (AU)\*** |  |  |  |  |
| grey matter | 48.5 ± 18.9 [44.7] | 30.6 ± 33.8 [19.9] | 8.3 ± 13.9 [2.6] | 2.7 ± 0.9 [2.5] |
| white matter | 28.8 ± 19.2 [21.4] | 13.8 ± 16.9 [2.7] | 6.2 ± 11.3 [0.8] | 2.9 ± 0.7 [2.9] |
| **VCAM-1 signal (AU)\*** |  |  |  |  |
| grey matter | 2.8 ± 0.4 [2.7] | 2.4 ± 1.3 [2.3] | 3.3 ± 1.3 [2.6] | 7.2 ± 2.9 [6.3] |
| white matter | 3.0 ± 0.8 [2.8] | 3.0 ± 1.2 [3.7] | 4.4 ± 1.8 [4.2] | 6.9 ± 2.1 [7.1] |

Linear mixed-effects models assessed per-field of view averaged vascular diameter and log-transformed per-field of view averaged vascular density, PECAM-1, PDGFRβ, and VCAM-1 signal using the lme4 R package[3], with cSVD group and tissue compartment (grey vs white matter) included as fixed-effect covariates. A case random effect was included to account for multiple non-independent measurements within each case, and models were formulated as follows:

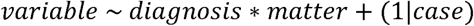

Here, variable was replaced with vascular diameter, log vascular density, log PECAM-1 signal, log PDGFRβ signal, and log VCAM-1 signal. ANOVA followed by estimated marginal means assessed the significance of group differences using the car and emmeans R package[29].

Gaussian mixtures modelling approach was used to identify potential patterns across the three markers PECAM-1, PDGFRβ, and VCAM-1. Log-transformed measures were scaled then processed with the mclust R package. Silhouette plots and PERMANOVAs were used to evaluate how well the diagnosis and markers-based clustering separated the cases. Statistics on the markers-based clusters were performed similarly to those based on the diagnostic groups with linear mixed-effect models and are presented in **Supplementary Figure 1** and **Table 3**.

**Table 3.** Quantitative vascular metrics by marker-based cluster. All metrics are reported as mean ± standard deviation (SD) [median]. Abbreviations: AU, arbitrary units; PECAM-1, platelet endothelial cell adhesion molecule-1; PDGFRβ, platelet-derived growth factor receptor-β; VCAM-1, vascular cell adhesion molecule-1. *, values multiplied by 10^3^ for display.

|  | Cluster 1 | Cluster 2 | Cluster 3 |
| --- | --- | --- | --- |
| N | 8 | 6 | 6 |
| **Vascular density (mm/mm<sup>3</sup>)** |  |  |  |
| grey matter | 880.4 ± 287.6 [816.5] | 1188.3 ± 359.3 [1048.4] | 1130.2 ± 459.5 [650.7] |
| white matter | 435.0 ± 108.3 [399.3] | 665.1 ± 106.3 [650.7] | 724.5 ± 143.1 [716.3] |
| **Average diameter (μm)** |  |  |  |
| grey matter | 8.7 ± 0.4 [8.7] | 8.5 ± 0.4 [8.6] | 8.5 ± 0.3 [8.5] |
| white matter | 9.7 ± 0.6 [9.7] | 9.6 ± 0.7 [9.7] | 9.8 ± 0.3 [9.8] |
| **PECAM-1 signal (AU)\*** |  |  |  |
| grey matter | 183.6 ± 22.7 [175.8] | 178.6 ± 28.8 [176.9] | 239.1 ± 21.9 [244.0] |
| white matter | 164.3 ± 26.6 [156.5] | 155.3 ± 24.2 [157.6] | 242.8 ± 35.9 [249.3] |
| **PDGFRβ signal (AU)\*** |  |  |  |
| grey matter | 50.4 ± 20.1 [44.7] | 5.2 ± 7.3 [2.7] | 2.7 ± 0.8 [2.5] |
| white matter | 29.0 ± 15.3 [24.2] | 1.5 ± 1.0 [1.3] | 2.9 ± 0.6 [2.9] |
| **VCAM-1 signal (AU)\*** |  |  |  |
| grey matter | 2.7 ± 0.4 [2.6] | 3.1 ± 1.7 [3.2] | 6.5 ± 3.2 [5.8] |
| white matter | 3.1 ± 0.8 [3.0] | 3.3 ± 1.3 [3.9] | 6.9 ± 1.9 [7.3] |

## Results

### Demographic comparisons

There were no differences in sex (5/15 females/males, p=1.00), post-mortem interval (p=0.30), hypercholesterolemia (p=1.00), diabetes (p=0.56), smoking status (p=0.32), or heart disease (p=0.92) between no SVD controls and increasing cSVD severity groups (**Table 1**). However, age at death increased with cSVD severity (no SVD controls: 32.6 ± 8.1 years; mild cSVD: 59.2 ± 18.4 years; moderate cSVD: 75.4 ± 7.7 years; severe cSVD: 83.2 ± 8.8 years; p=0.0043), consistent with the known age association of cSVD. Hypertension, one of the strongest risk factors for cSVD, was also found to increase in prevalence across disease groups (p=0.0075).

### Microvascular density, but not diameter, correlates with cSVD severity and age at death

Analyses were restricted to microvessels (mean diameter: 9.1 ± 2.8 μm), representing the most abundant vascular bed in brain tissue. Marked morphological differences were observed between tissue compartments, consistent with previous observations. In the no SVD control group, white matter vessels were 51% less dense (**Figure 2A, B, C**) and 14% wider (**Figure 2A, B, E**) than grey matter vessels.

**Figure 2.**
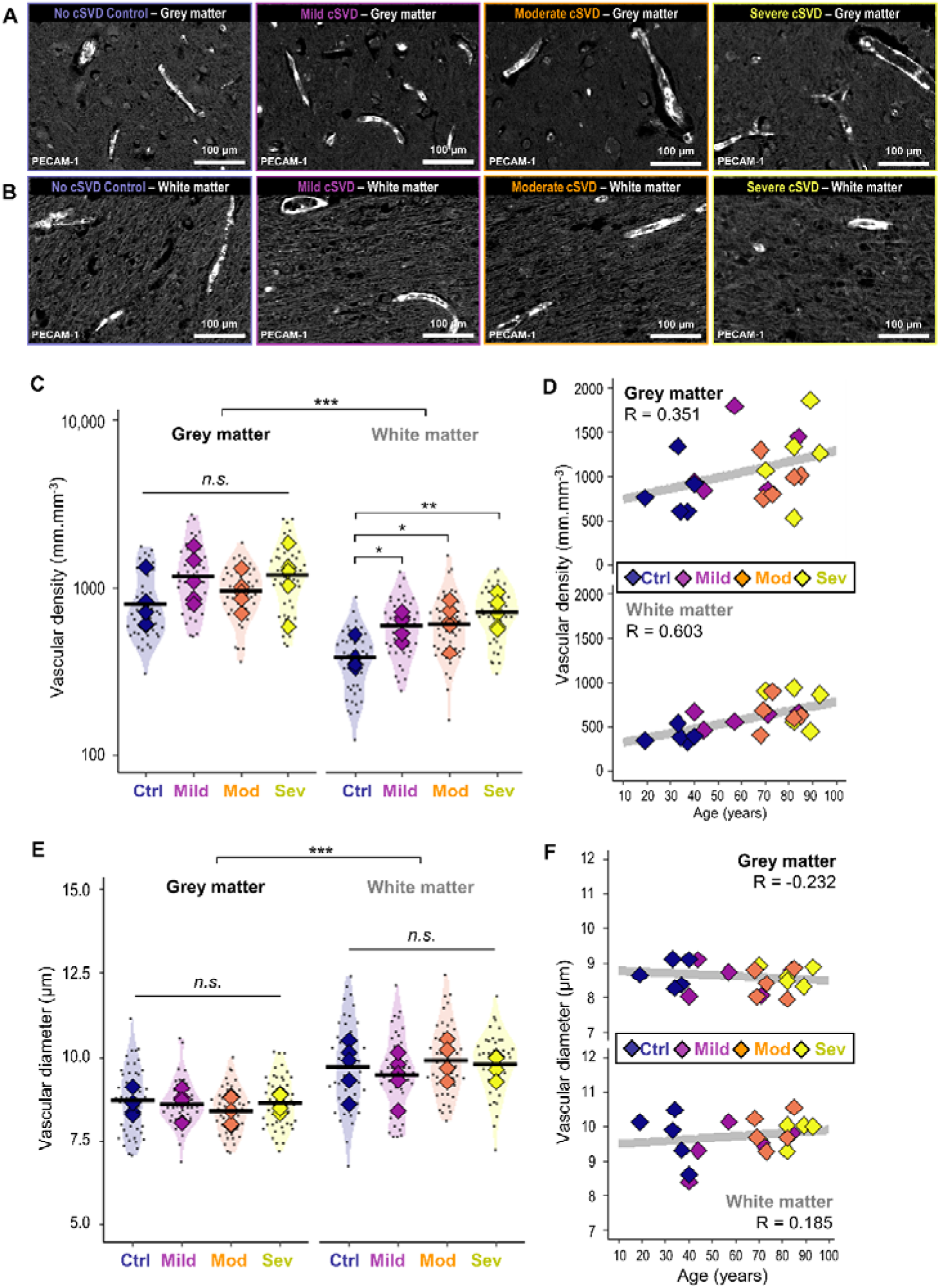
Cortical microvascular morphology across cSVD severity. (**A**, **B**) Representative PECAM-1 pseudo-fluorescent images in grey (A) and white (B) matter across cSVD severity groups (control, mild, moderate, severe). (**C**) Vascular density (vascular length per tissue volume) by group in grey and white matter. (**D**) Vascular density plotted against age at death. (**E**) Mean microvascular diameter by group in grey and white matter. (**F**) Microvascular diameter plotted against age at death. Diamond-shaped data points represent individual cases, whereas smaller dots represent analysed images. Scale bars: 100 μm. Linear mixed-effects modelling was used to account for case-level variability, with tissue compartment (grey/white matter) and images included as hierarchical factors. ANOVA on the fitted model identified significant differences between grey and white matter for vascular density and vascular diameter. Post-hoc comparisons were performed using estimated marginal means (emmeans), revealing significant differences in vascular density between cSVD groups in white matter (n.s., not significant; *, p-value < 0.05; **, p-value < 0.01; ***, p-value < 0.001). Pearson’s correlation coefficients (R) are shown in panels D and F. Abbreviations: cSVD, cerebral small vessel disease; Ctrl, no cSVD control; Mild, mild cSVD; Mod, moderate cSVD; Sev, severe cSVD; PECAM-1, platelet endothelial cell adhesion molecule-1.

While vascular density showed a trend towards increasing with cSVD status in both tissue compartments, it was statistically significant only in white matter, where the density of vessels doubled in severe cSVD cases compared to no cSVD controls (**Figure 2C**, p=0.0017) and increased with age (**Figure 2D**, Pearson’s R=0.603). Conversely, microvascular diameter did not change significantly across cSVD severity groups (**Figure 2E**) or with age (**Figure 2F**).

### VCAM-1 and PECAM-1 expression are markedly increased in the cortex of cases with severe cSVD

Representative images show low PECAM-1 and VCAM-1 signal in no cSVD controls and increased signal in severe disease (**Figure 3A, B**). PECAM-1 signal increased significantly with disease severity, displaying the highest value in the severe cSVD group, being 41% and 67% higher than in no cSVD controls in the grey and white matter, respectively (p=0.023 and 0.0002, respectively; **Figure 3C**). VCAM-1, which is typically expressed at low levels in capillaries, followed a similar trend, with its signal more than doubling (134% increase, p=0.0052 in grey matter; 155% increase, p=0.012 in white matter; **Figure 3G**).

**Figure 3.**
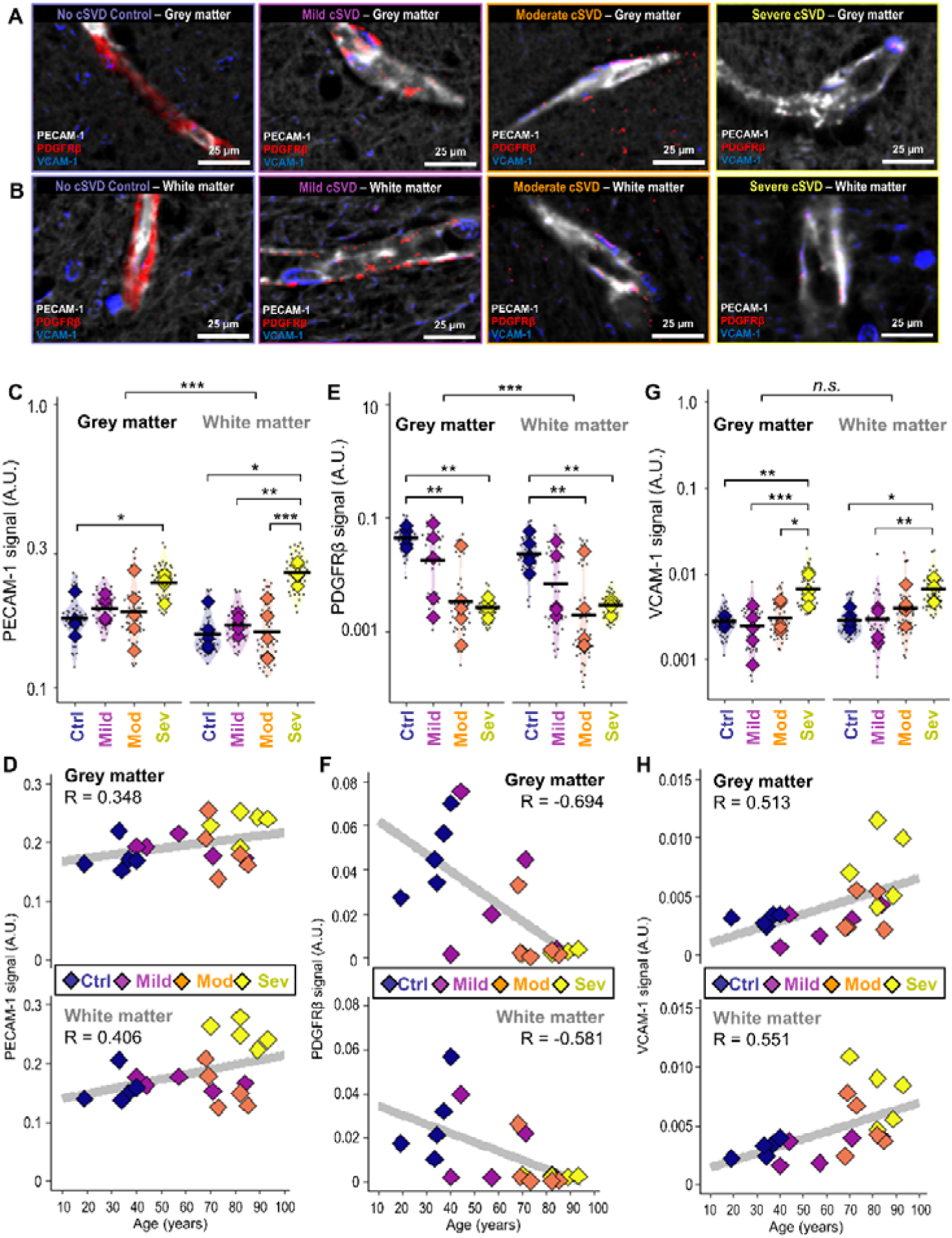
PECAM-1, PDGFRβ, and VCAM-1 signals across cSVD severity. (**A**, **B**) Representative microvessel close-ups in grey (A) and white (B) matter showing PECAM-1 (grey), PDGFRβ (red), and VCAM-1 (blue) across cSVD severity groups (control, mild, moderate, severe). (**C**) PECAM-1 signal by group in grey and white matter. (**D**) PECAM-1 signal plotted against age at death. (**E**) PDGFRβ signal by group in grey and white matter. (**F**) PDGFRβ signal plotted against age at death. (**G**) VCAM-1 signal by group in grey and white matter. (**H**) VCAM-1 signal plotted against age at death. Diamond-shaped data points represent individual cases, whereas smaller dots represent analysed images. Scale bars: 25 μm. Linear mixed-effects modelling was used to account for case-level variability, with tissue compartment (grey/white matter) and images included as hierarchical factors. ANOVA on the fitted model identified significant differences between grey and white matter for PECAM-1 and PDGFRβ signals. Post-hoc comparisons were performed using estimated marginal means (emmeans) to test differences between cSVD severity groups (n.s., not significant; *, p-value < 0.05; **, p-value < 0.01; ***, p-value < 0.001). Pearson’s correlation coefficients (R) are shown in panels D, F, and H. Abbreviations: AU, arbitrary units; cSVD, cerebral small vessel disease; Ctrl, no cSVD control; Mild, mild cSVD; Mod, moderate cSVD; Sev, severe cSVD; PECAM-1, platelet endothelial cell adhesion molecule-1; PDGFRβ, platelet-derived growth factor receptor-β; VCAM-1, vascular cell adhesion molecule-1.

This increase may reflect endothelial activation associated with ageing-related systemic inflammation (“inflamm-ageing”) or disease-specific vascular alterations. However, these changes did not show a clear linear relationship with either disease group or age (Pearson’s R=0.348 and 0.513 in grey matter, and 0.406 and 0.551 in white matter for PECAM-1 and VCAM-1, respectively; **Figure 3D, H**).

### Loss of PDGFR**β** signal in the cortex follows a bimodal distribution

PDGFRβ signal significantly decreased with increasing cSVD severity (**Figure 3A**), demonstrating an almost complete loss of pericytes in the most severely affected cases compared to no cSVD controls (94% loss, p=0.0038 in grey matter; 87% loss, p=0.0327 in white matter; **Figure 3E**). PDGFRβ signal was 52% lower in white matter than in grey matter in no cSVD controls, further emphasising the vulnerability of white matter.

Despite this difference between the tissues, PDGFRβ signal showed a strong negative correlation with both disease severity and age in grey (Pearson’s R=-0.694) and white (Pearson’s R=-0.581) matter (**Figure 3F**). However, the concept of a linear loss of pericyte coverage is challenged by the marked dichotomy observed within cases in the intermediate severity groups. Two cases from the mild group and one case from the moderate group displayed levels of pericyte coverage comparable to no cSVD controls, whereas the other cases showed an almost complete absence of PDGFRβ signal, like those from the severe cSVD group.

### Gaussian mixture clustering identifies three clusters based on marker signals

As visual inspection of the marker signal suggested the presence of underlying higher-order structure (**Figure 4A, B**), we performed a Gaussian mixture clustering analysis, with a gap statistic indicating three clusters, to reveal data stratification. Three clusters were obtained by this method, designated cluster 1 (C1), cluster 2 (C2), and cluster 3 (C3) in subsequent analyses (**Figure 4C**).

**Figure 4.**
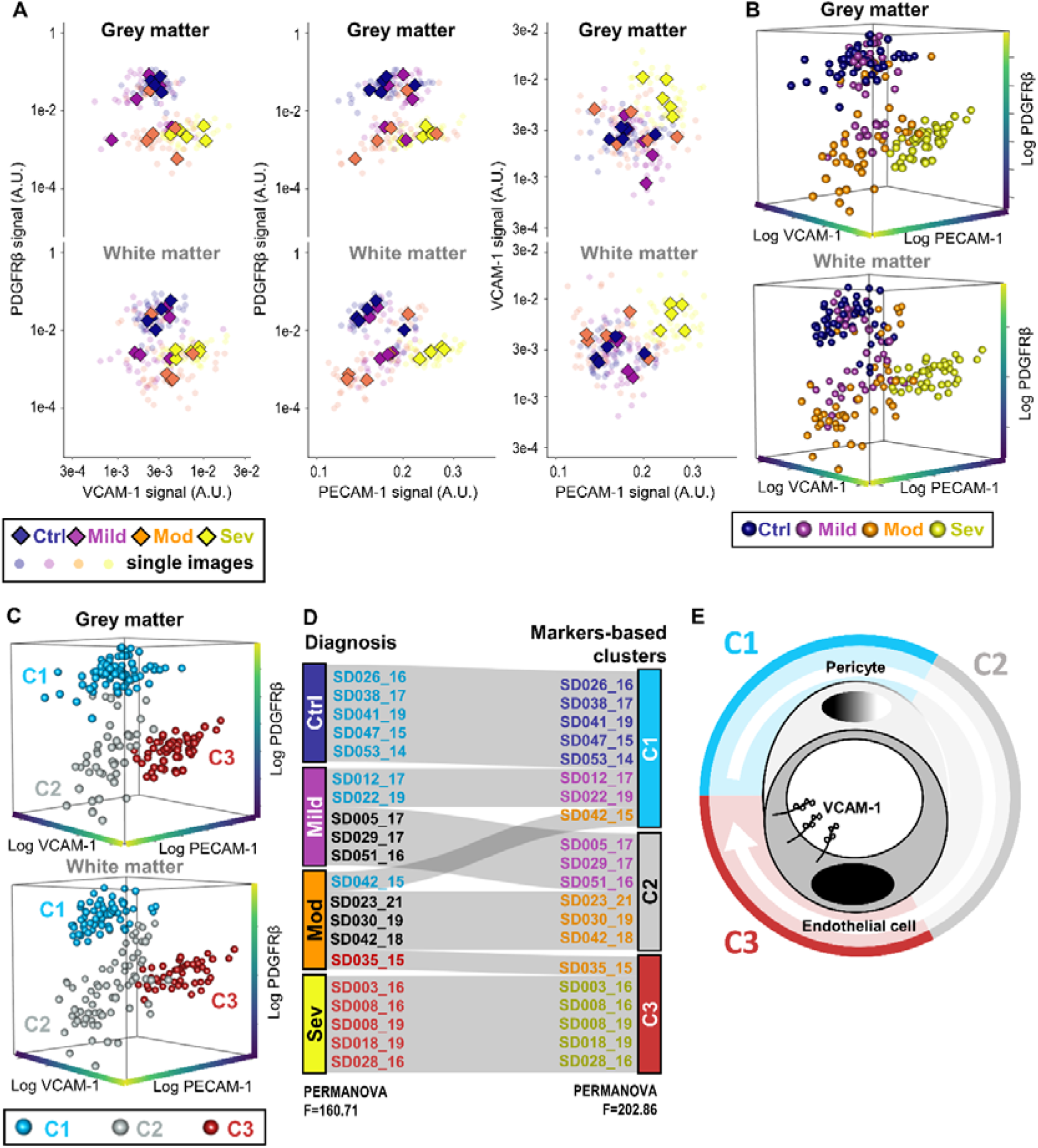
Marker-based clustering provides insights into disease progression. (**A**) Pairwise scatterplots between log-transformed PECAM-1, PDGFRβ, and VCAM-1 signals. Diamond-shaped data points represent individual cases, whereas smaller dots represent analysed images. (**B**) 3D scatterplot of log-transformed PECAM-1, PDGFRβ and VCAM-1 signals; colours indicate diagnostic group and points represent analysed images. (**C**) 3D scatterplot of the same markers coloured according to marker-based clusters identified by Gaussian mixture modelling. (**D**) Distribution of cases from diagnostic categories (left) to marker-based clusters (right). PERMANOVA F-statistics were computed using the pool of analysed images to compare clustering strength by diagnostic group versus marker-defined clusters. (**E**) Conceptual model of disease progression inferred from marker-based clusters. In cluster 1 (mostly no cSVD controls), pericyte coverage is preserved. In cluster 2 (mixed mild and moderate cSVD), pericyte coverage is reduced or lost. In cluster 3 (mostly severe cSVD), pericyte coverage is absent and endothelial cell adhesion molecules are strongly expressed. Abbreviations: AU, arbitrary units; C1, cluster 1; C2, cluster 2; C3, cluster3; cSVD, cerebral small vessel disease; Ctrl, no cSVD control; Mild, mild cSVD; Mod, moderate cSVD; Sev, severe cSVD; PECAM-1, platelet endothelial cell adhesion molecule-1; PDGFRβ, platelet-derived growth factor receptor-β; VCAM-1, vascular cell adhesion molecule-1.

C1 was characterised by high PDGFRβ and low PECAM-1 and VCAM-1, C2 by low PDGFRβ with low PECAM-1 and VCAM-1, and C3 by low PDGFRβ combined with high PECAM-1 and VCAM-1 expression. These differences were statistically significant (**Supplementary Figure 1**), and a higher PERMANOVA F-statistic was associated with the three clusters than with diagnostic alone (clusters: F=202.86; diagnosis: F=160.71), suggesting better separation of cases based on molecular marker profiles.

Intriguingly, the marker-based clustering largely recapitulated clinical diagnosis, with some notable deviations. C1 contained all no cSVD control cases, but also two cases from the mild group and one from the moderate group. C2 contained three cases from the mild group and three cases from the moderate group. C3 contained all severe cases, plus one from the moderate group (**Figure 4D**).

This distribution, which broadly parallels increasing disease severity, suggests that pericyte loss precedes endothelial activation, with vascular adhesion molecule upregulation emerging at later stages of disease progression (**Figure 4E**).

## Discussion

We applied an automated vascular segmentation and analysis pipeline[10] to multispectral imaging of cortical brain samples in a cohort of 20 cases chosen to represent increasing cSVD severity, and quantified the vascular morphology and molecular markers associated with pericyte coverage and endothelial activation in relation to disease severity. In this cohort of 20 cases from Brodmann area 4, we identify two key findings in the sequence of vascular degeneration in cSVD: (1) PDGFRβ signal loss occurs prior to the expression of endothelial cell adhesion molecules (CAMs), and (2) the bimodality of PDGFRβ signal suggests that pericyte coverage loss may occur abruptly rather than through a gradual, smooth decline.

In contrast to previous reports showing age- and disease-related capillary density losses[8, 51], our study observed an increase in the capillary network density with worse SVD. However, we focused on superficial layers of one area of frontal cortex, while the heaviest burden of cSVD-related vascular remodelling is usually reported in deeper regions. White matter included here was proximate to the cortex, therefore perhaps better represents compensation than deep tissues. A preclinical study investigating capillary densities in superficial cortical layers of aged mice found a slight increase in density compared to younger animals[5], and a clinical study in AD cases revealed generally preserved vascular structure and even higher capillary density[16]. Capillary density increases have been discussed as evidence of endothelial remodelling in response to tissue hypoxia[33]. Our findings may therefore reflect a compensatory adaptation to underlying hypoxia or hypoperfusion in deeper regions.

Pericytes are thought to be key stabilisers of endothelial proliferation and vascular homeostasis in health[30], and our concomitant finding of PDGFRβ signal loss with worse SVD further supports this role. White and grey matter also exhibited significant differences in capillary morphology, as previously reported[18]. The lower density observed in the white matter further emphasises the vulnerability of this region to vascular-related damage. Although capillaries were slightly larger in white matter than in grey matter, we did not detect any significant change in the average diameter of capillaries across cSVD severity groups.

The overall trends in the vascular markers investigated in this study are broadly consistent with previous reports. CAMs mediate cell-cell interactions, including between endothelial cells and circulating immune cells, and an increase in their expression with disease burden[15, 28] is not surprising in cSVD due to its major inflammatory component. Pericytes are major regulators of the vascular basement membrane and endothelial stability, extending processes that form extensive networks around endothelial cells[27]. The two cell types share intricate relationships through gap junctions, peg-and-socket contacts, and paracrine communication, whose intensity is thought to be reflected by the relative area of contact between them[40]. Pericyte coverage, here reflected by PDGFRβ signal, decreases with disease severity, supporting previous reports of pericyte loss in cSVD and other neurovascular dementias[14]. Pericyte loss also promotes white blood cell stalling by enhancing CAMs expression in endothelial cells [11].

Our high-resolution analysis revealed patterns inconsistent with a simple linear relationship between disease severity and vascular marker expression. PDGFRβ signal showed an apparent bimodal distribution across cases, while PECAM-1 and VCAM-1 signals increased primarily in the severe cSVD group. These findings suggest that pericyte alterations and endothelial activation may not progress in parallel, prompting further exploration of underlying patterns within the cohort.

As an initial approach to explore this potential complexity, we used Gaussian mixture clustering and identified three clusters among the 20 cases distributed across four diagnostic severities of cSVD burden based on pathologist-determined microvessel sclerosis and related features. Cluster 1 (C1) was characterised by low CAM expression and high pericyte coverage, largely overlapping with young no cSVD controls. Cluster 2 (C2) displayed pericyte coverage loss while CAM expression remained low and grouped cases in the intermediate ‘mild’ and ‘moderate’ diagnostic categories. Cluster 3 (C3) presented both loss of pericyte coverage and the highest levels of endothelial CAM expression. While interpretation of these results is limited by the sample size and the absence of age-matched controls across cSVD diagnosis, these findings motivate further investigations into the temporal course of microvascular alterations in cSVD.

Although this is a cross-sectional study in post-mortem brain, the gradation of severity within the marker-specific clusters suggests a sequential course of events in which pericyte coverage loss precedes CAM expression. This observation contrasts with our initial hypothesis that an inverse linear correlation would be observed between these markers. Instead, the clustering analysis suggests that pericyte depletion may persist for extended periods before endothelial activation becomes evident. Pericytes are known stabilisers of endothelial cells[27, 30], and no case was observed with both high pericyte coverage and high CAM expression. This observation raises the possibility of a therapeutic window in which restoration of pericyte function could occur before more extensive microvascular damage develops.

The bimodal distribution of PDGFRβ is another intriguing observation from this study. While the absence of intermediate degrees in PDGFRβ signal loss may partly reflect sampling limitations, it nevertheless suggests that pericyte coverage loss may not follow a linear decrease with age or cSVD burden. In this study, we propose that this distribution may reveal the potentially abrupt nature of pericyte coverage loss in cSVD. Previous studies reported decreases in pericyte number that appeared more linear with respect to case age[14, 19]. However, due to the imaging constraints associated with thin tissue sections, we could not count individual cells. This apparent discrepancy could be explained by the ability of pericytes to extend their processes to maintain endothelial coverage when neighbouring cells are lost[6]. In this scenario, pericyte numbers may decline while coverage remains relatively preserved until a critical threshold is reached, at which point coverage decreases abruptly. Given the central role of pericytes in stabilising capillaries, this transition may represent an important turning point in disease progression.

There are several important limitations to this study. The relatively small cohort size (n=20) and use of one tissue block from one mainly cortical brain region represent limitations; however, the absence of age-matched controls is a greater concern, as it partially confounds the distinction between cSVD-related changes and those associated with ageing. Importantly, identifying truly age-matched control individuals without some degree of WMH or microvascular pathology is increasingly difficult in older populations, which complicates the interpretation of vascular ageing in human cohorts. The gender balance is also limited, with only 5 of 20 participants being female, which may affect the generalisability of our findings. Although cSVD increases with age, this distribution may still introduce bias in the interpretation of vascular alterations, and sex differences in clinical presentations have been reported[26]. Additionally, despite the diagnosis of mild-to-moderate cSVD, three cases showed relatively high pericyte coverage, possibly reflecting regional resilience of the vasculature within the analysed cortical region. Finally, the apparent time gap between pericyte coverage loss and CAM expression may reflect additional mechanisms of vascular disruption that were not investigated in this study. Future studies should examine other brain regions in larger cohorts and ideally include age- and sex-matched non-cSVD controls. In addition, the contribution of other NGVU cell types, including astrocytes, microglia, and oligodendroglial populations, warrants investigation, as these cells are likely to interact with pericytes and endothelial cells to modulate microvascular dysfunction in cSVD.

In conclusion, this study provides new insights into the sequence of vascular alterations in cSVD. We propose that pericyte loss may represent an earlier marker of disease than endothelial CAM expression, and that this loss may occur over a relatively short timescale. Our results suggest that a critical threshold may exist in pericyte coverage, prompting future studies to characterise in larger populations the functional and morphological transitions of the cerebral microvasculature throughout cSVD progression.

## Acknowledgements

We are grateful to the donors and their families for their invaluable contribution to this research. We thank the teams involved in the Lothian Birth Cohort, the LINCHPIN study, and the Edinburgh Sudden Death Brain Bank for their essential work in tissue collection, processing, and clinical characterisation.

## Contributions

A.M. conceived and designed the study. D.J.G., C.M.Q., J.C.W., K.M.D., J.M.W., and C.S. contributed to data acquisition. A.C. and D.J.G. analysed the data. O.D. provided statistical advice. A.C. and A.M. wrote the first draft of the manuscript. All authors reviewed and approved the final manuscript.

---

**Supplementary Figure 1.**
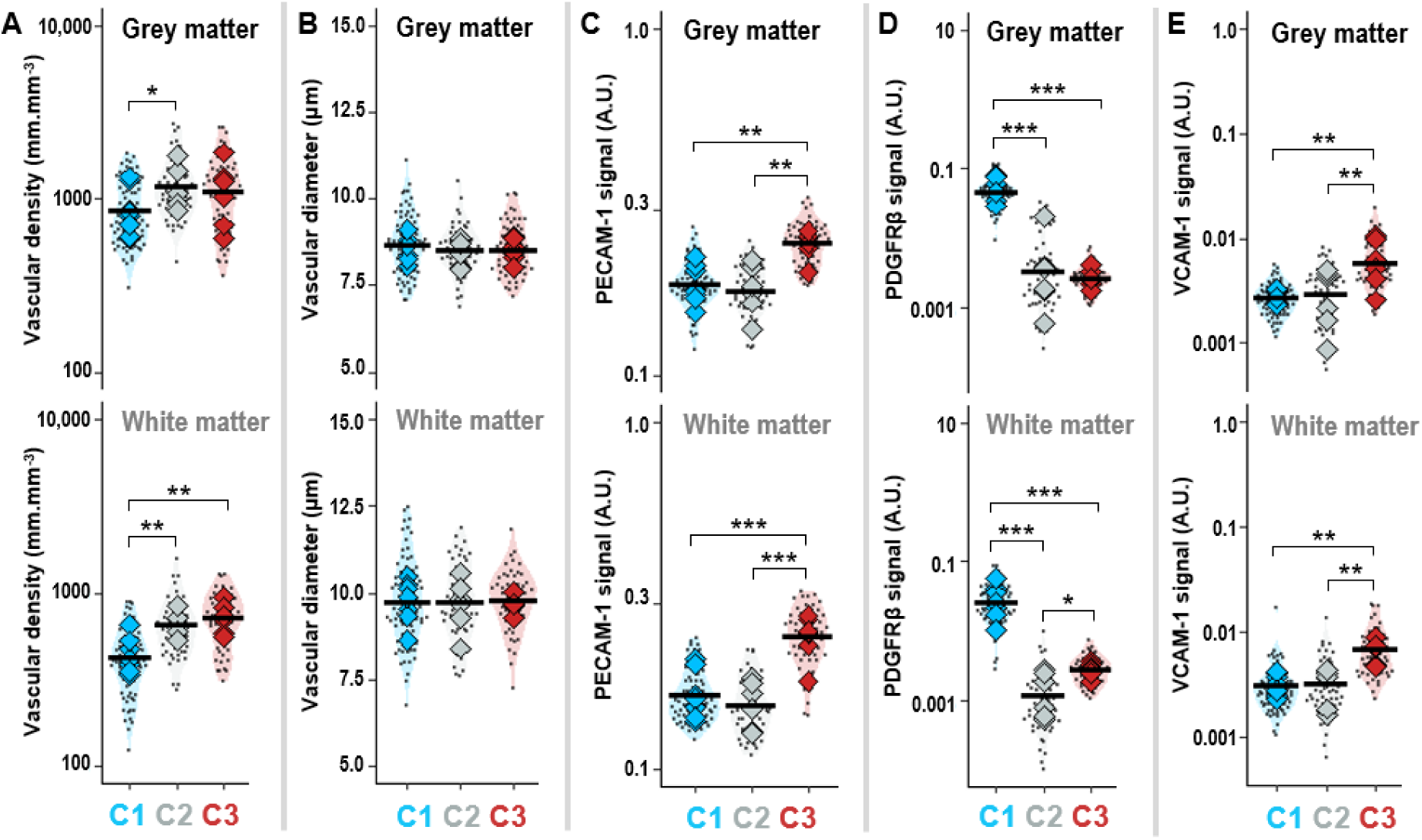
Vascular density, vascular diameter, PECAM-1, PDGFRβ, and VCAM-1 signals across marker-based clusters. Linear mixed-effects modelling was used to account for case-level variability, with tissue compartment (grey/white matter) and images included as hierarchical factors. (**A**) Vascular density, (**B**) vascular diameter, (**C**) PECAM-1 signal, (**D**) PDGFRβ signal, and (**E**) VCAM-1 signal. Post-hoc comparisons were performed using estimated marginal means (emmeans) and detected significant changes between clusters (*, p-value < 0.05; **, p-value < 0.01; ***, p-value < 0.001). Abbreviations: AU, arbitrary units; C1, cluster 1; C2, cluster 2; C3, cluster3; PECAM-1, platelet endothelial cell adhesion molecule-1; PDGFRβ, platelet-derived growth factor receptor-β; VCAM-1, vascular cell adhesion molecule-1.

